# Chitin-induced systemic disease resistance in rice requires both OsCERK1 and OsCEBiP and is mediated via perturbation of cell-wall biogenesis in leaves

**DOI:** 10.1101/2021.11.30.470685

**Authors:** Momoko Takagi, Kei Hotamori, Keigo Naito, Sumire Matsukawa, Mayumi Egusa, Yoko Nishizawa, Yuri Kanno, Mitsunori Seo, Shinsuke Ifuku, Akira Mine, Hironori Kaminaka

## Abstract

- Chitin is a well-known elicitor of disease resistance whose recognition by plants is crucial to perceive fungal infections. Chitin can induce both a local immune response and a systemic disease resistance when provided as a supplement in soils. Unlike local immune responses, how chitin-induced systemic disease resistance is deployed has not been studied in detail.
- In this study, we evaluated systemic disease resistance against the fungal pathogen *Bipolaris oryzae* by performing a transcriptome analysis and monitoring cell-wall composition in rice plants grown in chitin-supplemented soils. We also examined the local immune response to chitin by measuring the production of reactive oxygen species in leaves.
- Chitins induced both local immune response and systemic disease resistance with differing requirements for the receptors OsCERK1 and OsCEBiP. Transcriptome analysis suggested that a perturbation in cell-wall biogenesis is involved in the induction of systemic disease resistance, an idea which was supported by the induction of disease resistance by treatment with a cellulose biosynthesis inhibitor and alterations of cell-wall composition.
- These findings suggest that chitin-induced systemic disease resistance in rice is caused by a perturbation of cell-wall biogenesis in leaves through long-distance signalling after recognition of chitins by OsCERK1 and OsCEBiP.

## Introduction

Plants have developed two types of defence mechanisms, local immune response and systemic resistance, which are used to counteract threats from pathogens (Sun & Zhang, 2021). A local immune response is first induced upon pathogen approach and infection. Plants recognize microbe- or pathogen-associated molecular patterns (MAMPs/PAMPs) via a suite of pattern recognition receptors (PRRs) that induce pattern-triggered immunity (PTI), which causes the production of reactive oxygen species (ROS) and activates the expression of *Pathogenesis-related* (*PR*) genes to defend against pathogen invasion (Bittel & Robatzek, 2007; Zipfel, 2008). However, pathogens counteract this initial defence barrier by secreting effector proteins into plant cells that disrupt PTI and allow infection to progress. In response, plants have evolved nucleotide-binding/leucine-rich repeat receptors (NLRs) to recognize pathogen effectors, which induce a robust defence response often accompanied by a localised hypersensitive response (HR) leading to cell death. This form of immunity is called effector-triggered immunity (ETI) (Jones & Dangl, 2006; Cui *et al*., 2015).

Local pathogen infection triggers systemic acquired resistance (SAR) that spreads to distant non-infected cells and is associated with salicylic acid (SA)–dependent gene expression and the biosynthesis of secondary metabolites (Hartmann & Zeier, 2019). For instance, the synthetic SA-analogue benzothiadiazole (BTH), a chemical activator of SAR, can induce systemic resistance in tobacco (*Nicotiana tabacum*), wheat (*Triticum aestivum*), Arabidopsis (*Arabidopsis thaliana*), and rice (*Oryza sativa*) (Görlach *et al*., 1996; Friedrich *et al*., 1996; Lawton *et al*., 1996; Shimono *et al*., 2007). However, not all microbes negatively affect plant growth, as chemical treatments or beneficial microbes in the root microbiome collectively called plant growth–promoting rhizobacteria (PGPR) and fungi (PGPF) can trigger induced systemic resistance (ISR), which is mediated by long-distance signalling (Pieterse *et al*., 2014). Unlike the SA-dependent SAR pathway, ISR results in systemic resistance via multiple signalling pathways involving the phytohormones SA, jasmonic acid (JA), and ethylene (ET) (Pieterse *et al*., 1998, 2014). SAR and ISR engage different mechanisms but are both considered to elicit defence priming (Pieterse *et al*., 2014; Mauch-Mani *et al*., 2017).

Chitin, a polymer of β-1,4-linked *N*-acetylglucosamine, is a component of the fungal cell wall and arthropod exoskeletons (Pillai *et al*., 2009; Sharp, 2013). Plants have PRRs that recognize chitin as a MAMP/PAMP and initiate PTI (Gong *et al*., 2020). In rice, OsCERK1 (chitin elicitor receptor kinase 1) and OsCEBiP (chitin elicitor-binding protein) are members of the protein families lysin motif (LysM)-containing receptor-like kinase (RLK) and receptor-like protein (RLP) without a kinase domain, respectively; they form a heterodimeric chitin receptor complex (Kaku *et al*., 2006; Shimizu *et al*., 2010; Hayafune *et al*., 2014). OsCEBiP is the major chitin-binding protein in rice cultured cells (Kouzai *et al*., 2014b), with two OsCEBiP molecules binding to one chitin oligomer (CO) longer than a hexamer (Hayafune *et al*., 2014). By contrast, OsCERK1 does not directly bind to CO (Shinya *et al*., 2012) but mediates chitin-induced PTI by binding to and phosphorylating downstream factors (Kawasaki *et al*., 2017). OsCERK1 forms a heterodimer with the LysM-RLK OsMYR1 (Myc factor receptor 1), which perceives short-chain COs secreted by arbuscular mycorrhizal (AM) fungi and competitively inhibits OsCEBiP-dependent immune signalling (He *et al*., 2019; Zhang *et al*., 2021). In Arabidopsis, the LysM-RLK AtCERK1 is required for CO perception (Miya *et al*., 2007; Wan *et al*., 2008) by forming a receptor complex with AtLYK4 (LysM-containing receptor-like kinase 4) and AtLYK5 in the chitin signalling pathways (Cao *et al*., 2014).

Since natural polymeric chitin is difficult to use due to its intractability and insolubility (Pillai *et al*., 2009), water-soluble chitin forms such as COs have mainly been used in studies of plant immunity. We developed a method to produce chitin nanofiber (CNF) from original chitin polymers by simple physical treatment of crustacean exoskeletons (Ifuku & Saimoto, 2012). CNF can homogeneously disperse even in water and can be used as a solution of polymeric chitin. We previously reported that CNF, as well as a mixture of COs, elicits ROS production in Arabidopsis and rice and that spraying leaves with either COs or CNF enhances disease resistance against both the fungal pathogen *Alternaria brassicicola* and the bacterial pathogen *Pseudomonas syringae* pv. *tomato* DC3000 in Arabidopsis (Egusa *et al*., 2015). Moreover, CNF supplementation of soils induced systemic disease resistance in Arabidopsis, cabbage (*Brassica oleracea* var. *capitata*), and strawberry (*Fragaria* sp.) (Parada *et al*., 2018). In addition, treatment of rice roots with a CO solution induced systemic disease resistance for a day (Tanabe *et al*., 2006). The induction of ISR by the ectomycorrhizal fungus *Laccaria bicolor* on the nonmycorrhizal plant Arabidopsis was dependent on JA signalling and SA biosynthesis and signalling, and AtCERK1 was necessary for the effect of systemic resistance (Vishwanathan *et al*., 2020). Thus, although ISR induced by chitin or via chitin recognition has been studied, our knowledge about the molecular basis underlying the induction of systemic disease resistance by chitin is lacking compared to our understanding of local immune responses to chitins.

In this study, we examined the local immune response and systemic disease resistance against *Bipolaris oryzae*, the causal agent of rice brown spot disease, by performing a transcriptome analysis of rice plants treated with chitins. We exposed plants to both oligomeric chitin COs and polymeric chitin CNF to test the possibility of differential effects on chitin-induced disease resistance. Both chitins elicited ROS production and induced systemic disease resistance in leaves. Transcriptome analysis demonstrated that cell-wall biogenesis- and cytokinin-related genes are downregulated as a systemic response induced by chitins. We validated these results with a cellulose biosynthesis inhibitor, by monitoring cell-wall composition and quantifying phytohormone levels. Knockout mutants for *OsCERK1* and *OsCEBiP* revealed that both LysM receptors are required for chitin-induced systemic disease resistance in response to *oryzae* but not for the local immune response in leaves.

## Materials and Methods

### Plant growth conditions

Unless otherwise stated, the Nipponbare cultivar of *Oryza sativa* L. (*japonica* group) was used as wild-type rice. *OsCEBiP* or *OsCERK1* transformant lines were generated in the *Oryza sativa* L. *japonica* ‘Nipponbare Kanto BL number 2’ background, which was previously described (Kouzai *et al*., 2014b,a). The knockout mutant and segregating wild-type siblings of *oscebip* line 169 and *oscerk1* lines 19 and 53 were used in this study. Rice seeds were soaked in distilled water (DW) for germination at 28°C for 3 or 4 days in the dark, and the germinated seeds were transplanted into sterilized culture soil (Bestmix No. 3; Nippon Rockwool, Japan) mixed with 0.1 or 0.01% (w/v) CO solution and CNF dispersion in DW at 1% (w/v) in magenta boxes (GA-7; Sigma-Aldrich, USA). Plants were grown in a growth cabinet (BiOTRON; NK-systems, Japan) under controlled conditions (28°C 14-h-light/25°C 10-h-dark cycles) and fertilised once a week with a 1:1000 HYPONeX (6-10-5; HYPONeX, Japan) solution. The COs (the mixture of DP [degree of polymerization] 2-6 chitin oligomers; NA-COS-Y; Yaizu Suisankagaku Industry, Japan) solution and CNF dispersion in water were prepared as previously reported (Kaminaka *et al*., 2020). The cell-wall biosynthesis inhibitor isoxaben (ISX; Santa Cruz Biotechnology, Germany) was resolved in dimethyl sulfoxide (DMSO), and a dilution in DW was sprayed onto leaves 5 h before sampling.

### ROS measurements

Fourth leaves from 3-week-old rice seedlings grown on soil without chitin supplementation were excised into nine leaf discs 0.5 mm in size and floated overnight at 22°C in a well filled with sterilized DW (sDW). COs or CNF elicitation solutions were prepared by suspending COs or CNF in sDW to a final concentration of 0.01% (w/v). Peroxidase from a horseradish root (HRP: Oriental Yeast, Japan) stock solution (500×HRP) and luminol L-012 (L-012; Wako, Tokyo, Japan) stock solution (20 mM) were prepared as previously described (Parada *et al*., 2018). Before elicitation, the sDW was carefully removed from each well without tissue damage or desiccation. The elicitation solution was immediately added to each well after removing sDW, and chemiluminescence was measured with a microplate reader (ARVO X3; PerkinElmer Japan, Japan) for 40 min.

### Pathogen inoculation test

*Bipolaris oryzae* D6 (Kihara & Kumagai, 1994) was cultured on potato dextrose agar plates for 1 week at 25°C in the dark. The conidial suspension was prepared to a titre of 1×10^5^ spores/mL in 0.25% (v/v) Tween 20. Fourth leaves from 3-week-old rice seedlings grown on normal or chitin-supplemented soils were detached and inoculated with a drop (5 µL) of spores on the leaf sheaths and incubated in the dark for 1 day and then in the light for 1 day at 25°C. Images of inoculated leaves were taken using a GT-S640 Scanner (EPSON, Japan), and each lesion diameter was measured by ImageJ (ver.1.53a).

### Phytohormone measurements

Approximately 500 mg of randomly selected leaves was excised from at least three individual 3-week-old rice seedlings grown on normal or chitin-supplemented soils. Samples were prepared with five technical replicates for each treatment. The leaves were placed in tubes and frozen in liquid nitrogen. The contents of each phytohormone were quantified using liquid chromatography–tandem mass spectrometry (LC-MS/MS) as previously described (Kanno *et al*., 2016).

### Fourier-transform infrared (FT-IR) spectroscopy

Alcohol-insoluble residue (AIR) was prepared from excised third or fourth leaves of 3-week-old rice seedlings grown on normal or chitin-supplemented soils, according to Bacete *et al*. (2017). AIR fractions were subjected to FT-IR spectroscopy using an FT-IR spectrophotometer equipped with an attenuated total reflectance accessory (Spectrum 65; PerkinElmer Japan, Japan). The FT-IR spectra were collected in the wavenumber range from 600 to 4,000 cm^−1^ with 16 scans, and the average values of three AIR fractions obtained from independent plants were used.

### Transcriptome deep sequencing (RNA-seq) and data analysis

Rice plants were grown as mentioned above except for the growth conditions (28°C 14-h-light/16°C 10-h-dark cycles). About 100 mg of randomly selected leaves or roots was 10 excised from at least three individual 3-week-old rice seedlings grown on normal or chitin-supplemented soils. Samples were prepared from three technical replicates for each treatment. The leaves or roots were placed inside tubes with 5-mm stainless beads, frozen in liquid nitrogen, and pulverised for 30 s using ShakeMan 6 (Bio Medical Science, Japan). LBB solution [1 M LiCl, 100 mM Tris-HCl (pH 7.5), 1% SDS, 10 mM EDTA, 0.015% Antifoam A, 5 mM DTT, and 71.5 mM 2-ME, DNase/RNase-free water] was added to the samples and completely dissolved by vortexing. All samples were incubated for at least 5 min at room temperature with occasional inverting and mixing. After centrifugation at 20,630 *g* for 10 min at room temperature, the supernatant was transferred to new tubes and stored at –80°C. Sequencing libraries were produced according to the BrAD-seq protocol (Ichihashi *et al*., 2018). Sequencing was performed on a HiseqX instrument (Illumina) by Macrogen Japan. Raw reads were checked for quality, and adaptor sequences were trimmed using fastp (Chen *et al*., 2018). The resulting clean reads were mapped to the reference rice genome (MSU Rice Genome Annotation Project ver. 7.0; http://rice.plantbiology.msu.edu/) using STAR (Dobin *et al*., 2013), and reads were counted by featureCounts with the package Subread (Liao *et al*., 2014). Results of data analysis are summarised in Table **S1**. The expression profiles were obtained by comparing control and chitin-treated plants using EdgeR in the R package (Robinson *et al*., 2010) with a trimmed mean of M values for normalisation. The list of differentially expressed genes (DEGs) was based on a false discovery rate (FDR) < 0.05. Venn diagrams and heatmaps were prepared at the Bioinformatics and Evolutionary Genomics webpage (http://bioinformatics.psb.ugent.be/webtools/Venn/) and ComplexHeatmap in R package (Gu *et al*., 2016), respectively. Gene Ontology (GO) enrichment analysis was conducted with the PANTHER (Mi *et al*., 2021) and REVIGO (Supek *et al*., 2011) tools, according 11 to Bonnot *et al*. (2019). Co-expression analysis was performed using the ShinyGO (ver.0.61) website (http://bioinformatics.sdstate.edu/go/; Ge *et al*., 2020).

## Results

### Both oligomeric and polymeric forms of chitin induce a local immune response in rice leaves

COs induce ROS production, a typical response of PTI, in Arabidopsis and rice (Kaku *et al*., 2006; Miya *et al*., 2007). Furthermore, polymeric chitin in the form of CNF induces ROS production in Arabidopsis seedlings, rice cultured cells, and cabbage and strawberry leaf discs (Egusa *et al*., 2015; Parada *et al*., 2018). We confirmed that both COs and CNF induce ROS production in rice leaf discs (Fig. **1a**). We previously showed that CNF induces a more pronounced ROS production than COs (of DP2-6 and DP6) when treating Arabidopsis seedlings and rice cultured cells (Egusa *et al*., 2015); however, we observed the opposite result in rice leaves (Fig. **1a**). These results nevertheless indicated that both COs and CNF can induce a local immune response in rice leaves.

**Fig. 1.**
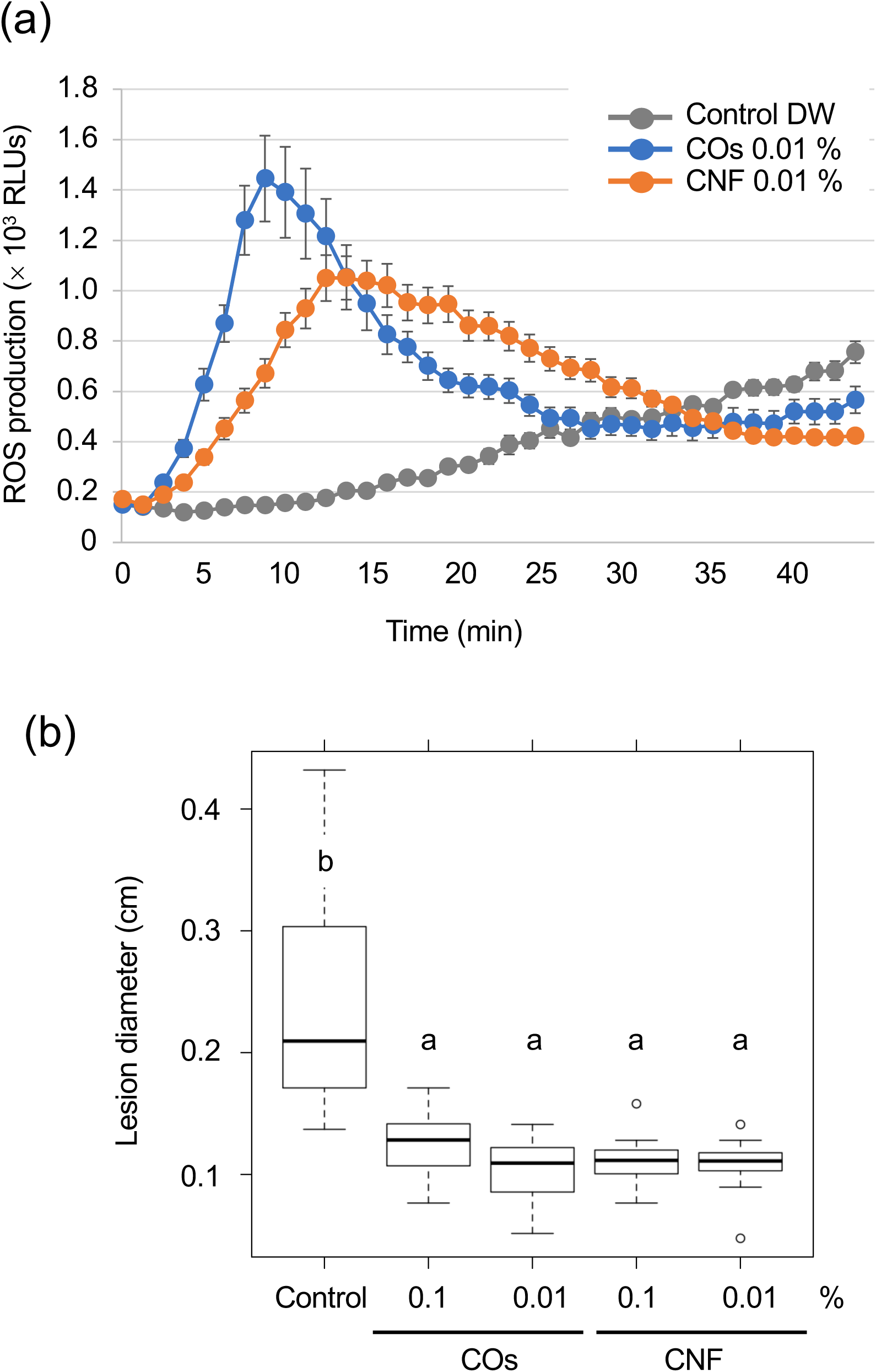
Chitins induce a local immune response and systemic disease resistance in rice. (a) Reactive oxygen species (ROS) production induced by chitins in rice leaves. Leaf discs were prepared from 3-week-old rice seedlings grown on soil and treated with distilled water (DW; Control), 0.01% (w/v) chitin oligomers (COs), or chitin nanofiber (CNF). ROS production triggered by chitin treatments was measured in relative luminescence units (RLUs) in the presence of the probe L-012 for 40 min. Representative data from three independent experiments are shown. Error bars, standard deviation (SD, *n* = 6). (b) Induced systemic disease resistance against *Bipolaris oryzae* by chitins, as measured by lesion diameter (in centimetres) of leaves 2 days after inoculation. Three-week-old rice seedlings grown on soil mixed with DW (Control) and 0.1% or 0.01% (w/v) COs and CNF were inoculated with *B. oryzae*. Representative results from three independent experiments are shown. Error bars, SD (*n* > 14). Different letters indicate significant differences by Tukey’s test (*p* < 0.05).

### Chitin supplementation of soils induces systemic disease resistance in rice leaves

We examined the level of systemic disease resistance in chitin-treated rice plants using the rice brown spot fungus *B. oryzae*. Supplementation of soils with COs and CNF solution/dispersion induced disease resistance compared to untreated control plants, as determined by the size of lesions on leaves; both chitin forms had comparable effects (Fig. **1b**). Mounting an immune response is often accompanied by growth inhibition, a trade-off between immunity and growth (Huot *et al*., 2014). Supplementation of soils with 0.1% (w/v) CNF hinders the development of cabbage and strawberry plants (Parada *et al*., 2018). However, 0.1% CNF added to soils did not affect leaf or stem growth in rice seedlings (Fig. **S1**). These results revealed that both COs and CNF can systemically induce disease resistance without compromising growth in rice.

### Transcriptome analysis of rice plants grown in soils supplemented with chitins

To explore the molecular mechanisms underlying the induction of systemic disease resistance by chitins, we performed an RNA-seq analysis of rice leaves and roots grown in soils mixed with COs or CNF. We identified 81 and 230 DEGs in CO- and CNF-treated rice leaves, respectively (FDR < 0.05; numbered in both MSU-DB and RAP-DB) compared to control leaves (Fig. **S2**; Tables **S2**, **S3**). Of these 297 non-redundant DEGs, only 14 genes were shared between CO and CNF treatments (Fig. **S2**). The 297 DEGs consisted of 157 upregulated (LogFC > 0) and 140 downregulated (LogFC < 0) genes by chitin treatments and showed similar trends in their expression patterns in CO- and CNF-treated leaves (Fig. **2a**). A GO enrichment analysis of DEGs indicated an enrichment for categories “regulation of protein serine/threonine phosphatase activity (GO:0080163)”, “glutathione metabolic process (GO:0006749)”, and “cellular modified amino acid metabolic process (GO:0006575)” among upregulated genes (Fig. **2b**), while downregulated genes were associated with “sulfate assimilation (GO:0000103)”, “response to cytokinin (GO:0009735)”, “cytokinesis (GO: 0000910)”, and “cell wall biogenesis (GO:0042546)” (Fig. **2c**). A co-expression analysis conducted using ShinyGO (Ge *et al*., 2020) determined that the expression of 41 genes upregulated by chitins was strongly and significantly (2.18 × 10^−48^) correlated with BTH-induced genes (Fig. **S3**; Table **1**; Shimono *et al*., 2007).

**Fig. 2.**
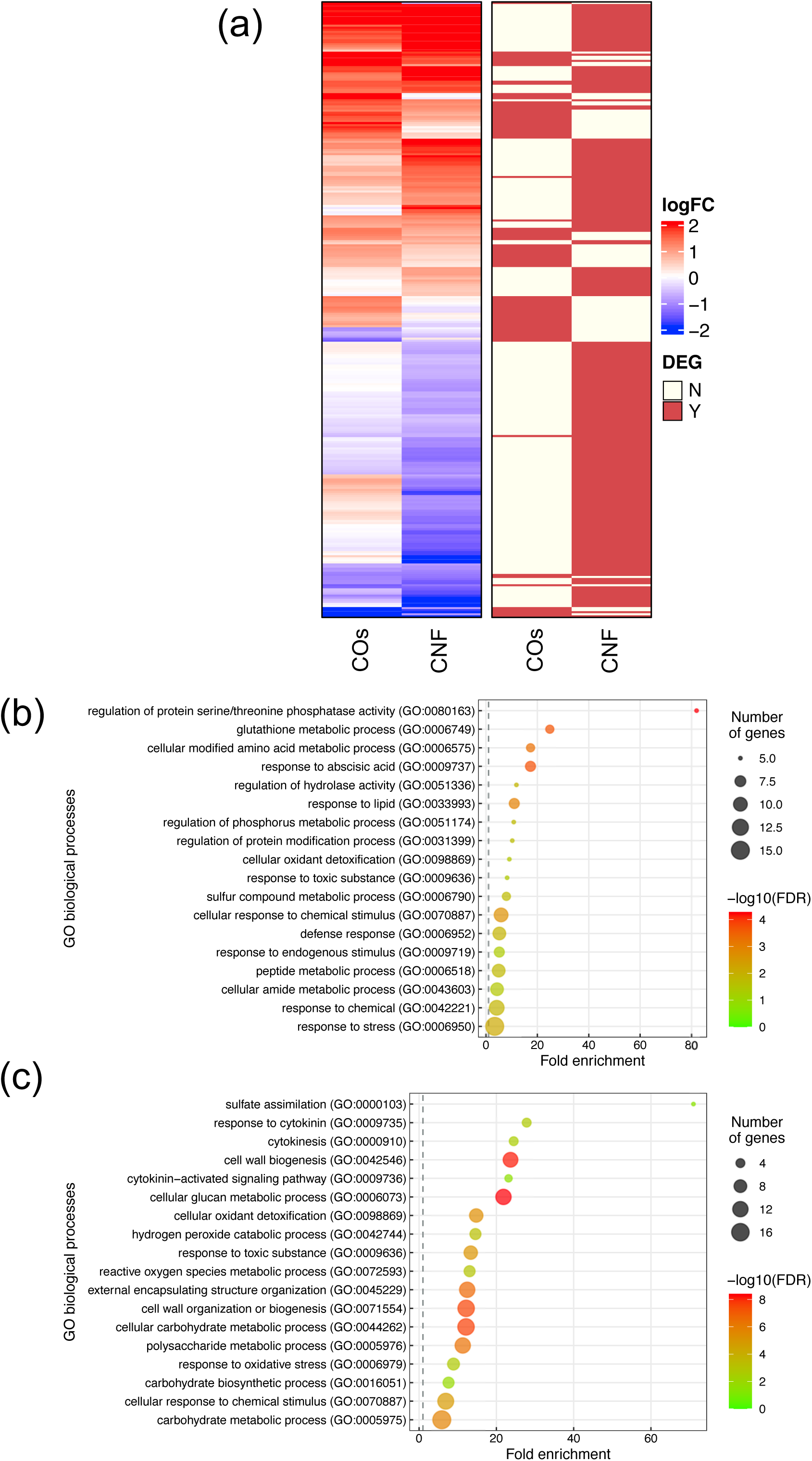
Transcriptome analysis of rice leaves grown on chitin-supplemented soils. (a) Heatmap representation of gene expression levels of differentially expressed genes (DEGs) in response to COs and CNF (left). LogFC is shown between −2 and 2, with outside values indicated as 2 or −2. Red, upregulated genes; blue, downregulated genes. DEGs in each treatment are indicated on the right. Red, DEG; ivory, not differentially expressed. (b, c) Results of Gene Ontology (GO) enrichment analysis summarised as plot data for upregulated DEGs in chitin-treated samples (b) and downregulated DEGs (c).

**Table 1.** Expression levels and annotation of genes upregulated by chitin treatment correlated with BTH-induced genes.

| Gene ID | COs LogFC | CNF LogFC | Annotation |
| --- | --- | --- | --- |
| Os01g0176000 | 0.632 | 1.487 | flavonol-3-*O*-glycoside-7-*O*-glucosyltransferase 1, putative, expressed |
| Os01g0510200 | 1.921 | 3.168 | expressed protein |
| Os01g0585200 | 0.883 | 1.784 | expressed protein |
| Os01g0627800 | 0.904 | 2.160 | cytochrome P450 72A1, putative, expressed |
| Os01g0638000 | 1.507 | 3.712 | anthocyanin 3-*O*-beta-glucosyltransferase, putative, expressed |
| Os01g0695800 | 0.767 | 1.242 | ABC transporter, ATP-binding protein, putative, expressed |
| Os01g0795200 | 2.290 | 3.552 | OsSub8 - Putative Subtilisin homologue, expressed |
| Os02g0726700 | 1.111 | 2.136 | helix-loop-helix DNA-binding domain containing protein, expressed |
| Os03g0235000 | 1.745 | 0.875 | peroxidase precursor, putative, expressed |
| Os03g0757600 | 1.590 | 2.651 | UDP-glucuronosyl and UDP-glucosyl transferase domain containing protein, expressed |
| Os04g0339400 | 2.810 | 4.817 | oxidoreductase, aldo/keto reductase family protein, putative, expressed |
| Os04g0373400 | 0.796 | 1.494 | MATE efflux family protein, putative, expressed |
| Os04g0581000 | 1.475 | 0.702 | naringenin,2-oxoglutarate 3-dioxygenase, putative, expressed |
| Os04g0627900 | 1.507 | 2.599 | expressed protein |
| Os05g0527000 | 1.658 | 2.435 | anthocyanidin 5,3-*O*-glucosyltransferase, putative, expressed |
| Os06g0493100 | 1.310 | 0.905 | RALFL28 - Rapid ALkalinization Factor RALF family protein precursor, expressed |
| Os07g0175600 | 1.549 | 0.882 | LTPL78 - Protease inhibitor/seed storage/LTP family protein precursor, expressed |
| Os07g0442900 | 2.977 | 3.307 | membrane associated DUF588 domain containing protein, putative, expressed |
| Os07g0605400 | 0.746 | 0.450 | EGG APPARATUS-1, putative, expressed |
| Os07g0677200 | 1.227 | 0.694 | peroxidase precursor, putative, expressed |
| Os08g0153900 | 2.204 | 0.956 | expressed protein |
| Os08g0155900 | 1.831 | 3.026 | expressed protein |
| Os08g0185900 | 1.146 | 1.798 | ubiquitin family protein, putative, expressed |
| Os08g0412700 | 1.173 | 1.989 | expressed protein |
| Os09g0492900 | 2.017 | 4.108 | expressed protein |
| Os10g0109600 | 2.748 | 1.908 | peroxidase precursor, putative, expressed |
| Os10g0527400 | 2.717 | 3.543 | glutathione *S*-transferase GSTU6, putative, expressed |
| Os10g0527800 | 3.119 | 3.641 | glutathione *S*-transferase, putative, expressed |
| Os10g0529500 | 1.311 | 2.201 | glutathione *S*-transferase GSTU6, putative, expressed |
| Os10g0535800 | 0.945 | 0.954 | uncharacterized Cys-rich domain containing protein, putative, expressed |
| Os10g0542900 | 2.095 | 1.632 | CHIT14 - Chitinase family protein precursor, expressed |
| Os10g0558700 | 0.932 | 1.340 | flavonol synthase/flavanone 3-hydroxylase, putative, expressed |
| Os11g0687100 | 2.968 | 1.771 | von Willebrand factor type A domain containing protein, putative, expressed |
| Os12g0268000 | 1.387 | 2.397 | cytochrome P450 71A1, putative, expressed |
| Os12g0555200 | 1.568 | 1.714 | pathogenesis-related Bet v I family protein, putative, expressed |
| Os12g0555500 | 0.732 | 0.319 | pathogenesis-related Bet v I family protein, putative, expressed |

We conducted a similar analysis on root samples (Fig. **S4**; Tables **S4**, **S5**). Roots exhibited a much smaller number of DEGs compared to that of leaves upon chitin treatment (Figs. **S2, S4**). GO enrichment analysis revealed that upregulated genes in response to COs and CNFs are involved in “cellular response to nitrate (GO:0071249)”, “nitrogen cycle metabolic process (GO:0071941)”, and “nitrate assimilation (GO:0042128)” (Fig. **S4c**). These results demonstrated that both COs and CNF induce the expression of genes involved in cytokinin signalling, cell-wall biogenesis, and defence priming in leaves, while the influence of chitin supplementation in soils was more limited in roots.

### Chitin supplementation of soils affects endogenous cytokinin levels and cell-wall composition in rice leaves

Phytohormones plays important roles in ISR (Pieterse *et al*., 2014; Hartmann & Zeier, 2019). We thus measured endogenous levels of phytohormones (auxin [IAA], gibberellins [GA_1_], abscisic acid [ABA], JA, jasmonyl isoleucine [JA-Ile], *trans*-zeatin [tZ], isopentyladenine [iP], and SA) in the leaves of rice seedlings grown on soils supplemented with chitins (Table **2**). Of all phytohormones tested, only the contents for the active cytokinin tZ significantly (9.71 × 10^−4^) decreased in CNF-treated samples compared to control seedlings. This finding was congruent with our RNA-seq analysis showing that the GO term “response to cytokinin” was enriched in chitin-suppressed genes in leaves (Fig. **S5**).

**Table 2.** Levels of endogenous phytohormones upon induction by chitins.

|  | Control | 0.1% COs | 0.1% CNF |
| --- | --- | --- | --- |
| IAA | 68.37±13.48 | 84.08±43.8 | 72.39±22.83 |
| GA<sub>1</sub> | 4.14±1.08 | 4.42±2.26 | 3.06±0.71 |
| ABA | 23.60±0.63 | 25.54±11.22 | 25.40±7.69 |
| JA | 55.55±48.20 | 29.02±18.93 | 16.08±8.66 |
| JA-Ile | 9.14±6.55 | 3.41±2.03 | 2.28±1.09 |
| tZ | 2.98±0.26 | 3.51±1.30 | ***1.45±0.55 |
| iP | 0.39±0.03 | 0.46±0.24 | 0.35±0.11 |
| SA | 48.53±11.00 | 49.19±18.64 | 47.55±16.04 |

\*\*\*, $p < 0.001$ , $n = 5$ ; IAA, GA<sub>1</sub>, ABA, JA, JA-Ile, tZ, iP, pg/mg; SA, ng/mg.

The plant cell wall offers a passive physical defence barrier to prevent pathogen access to plant cells; in agreement, alteration of cell-wall composition is associated with disease resistance (Bacete *et al*., 2018). Modification of cell-wall composition caused by 14 genetic inactivation or overexpression of cell-wall-related genes in Arabidopsis resulted in enhanced disease resistance or susceptibility against various pathogens (Bacete *et al*., 2018; Molina *et al*., 2021). Since the GO enrichment analysis suggested an alteration of cell-wall composition in leaves by chitin supplementation of soils (Fig. **2c**), we purified the cell-wall fraction from leaves of control and chitin-treated rice seedlings and measured its absorbance using FT-IR spectroscopy. We selected the wavenumber range of 800–1700 cm^−1^ as in Molina *et al*. (2021), which can be assigned to main cell-wall components (Alonso-Simón *et al*., 2011). As shown in Fig. **3a**, we observed different FT-IR spectra in seedlings grown on chitin-supplemented soils, indicating that chitin supplementation of soils results in an alteration of cell-wall composition in rice leaves.

**Fig. 3.**
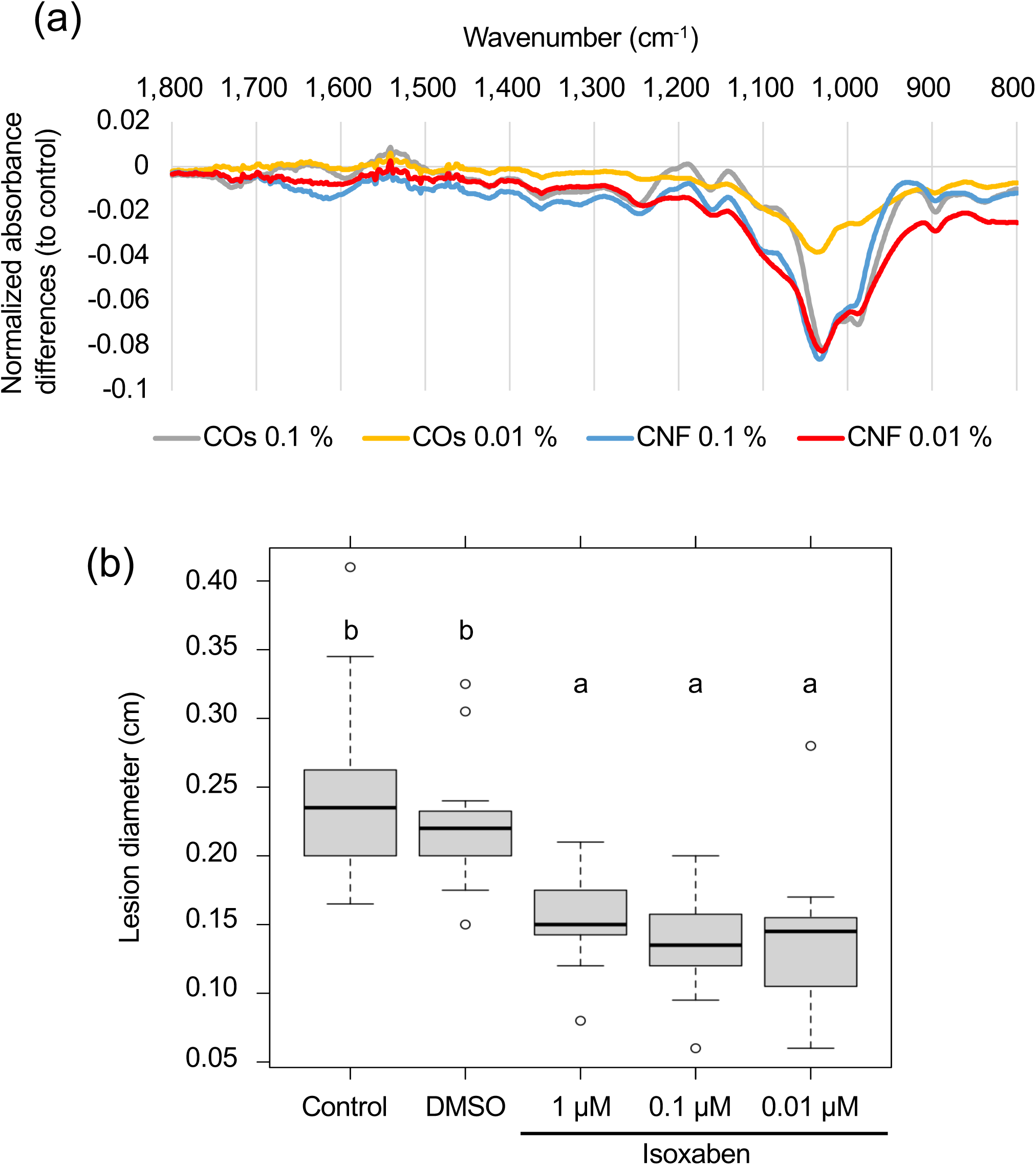
Perturbation of cell-wall biogenesis upon chitin treatments and induced systemic disease resistance in rice leaves. (a) Supplementation of soils with chitins systematically induced alterations of cell-wall composition in rice leaves. Each line represents the differential Fourier-transform infrared (FT-IR) spectra between control plants and seedlings grown on CO- or CNF-containing soils (*n* = 3). (b) A cellulose biosynthesis inhibitor induces disease resistance in rice leaves. The leaves of 3-week-old rice seedlings grown on soil were sprayed with control (DW), DMSO, or isoxaben (1 , 0.1 , and 0.01 µM, respectively) 5 h before *B. oryzae* inoculation. Lesion diameter (cm) of leaves 2 days after inoculation are shown. Representative results from three independent experiments are shown. Error bars, SD (*n* > 14). Different letters indicate significant differences by Tukey’s test (*p* < 0.05).

### Cellulose biosynthesis inhibition induces disease resistance in rice leaves

Defects in cellulose biosynthesis are associated with disease resistance against *Plectosphaerella cucumerina*, *Botrytis cinerea*, and *Ralstonia solanacearum* in Arabidopsis (Hernández-Blanco *et al*., 2007). However, an effect of cellulose biosynthesis inhibition on disease resistance has not been reported in rice. Accordingly, we examined disease resistance against *B. oryzae* in rice leaves treated with the cellulose biosynthesis inhibitor ISX (Heim *et al*., 1990; Tateno *et al*., 2016). ISX treatment significantly enhanced disease resistance against *B. oryzae*, compared to control and DMSO-treated seedlings (*p* < 0.005), indicating that, as in Arabidopsis, inhibition of cellulose biosynthesis increases resistance against pathogens in rice (Fig. **3b**).

### Both LysM receptors, OsCERK1 and OsCEBiP, are required to induce systemic disease resistance by chitins in rice leaves

To assess whether OsCERK1 or OsCEBiP contributes to the local immune response in chitin-treated rice seedlings, we quantified ROS production in their respective knockout mutants and corresponding wild-type segregants (Kouzai *et al*., 2014b,a), which we used as wild-type plants in the following experiments. ROS production by chitins was compromised in the *oscerk1* mutant compared to its wild-type siblings (Fig. **4a**). However, ROS production was comparable between the *oscebip* mutant and its wild-type siblings (Fig. **4b**). We also tested chitin-induced systemic disease resistance in all genotypes and observed that both wild-type siblings and wild-type plants exhibited a systemic induction of disease resistance against *B. oryzae* upon chitin treatments, whereas neither knockout mutant did (Fig. **5**). Taken together, these findings indicate that both OsCERK1 and OsCEBiP are required for chitin-induced systemic disease resistance in rice, but OsCEBiP did not appear to be essential for a local immune response in leaves.

**Fig. 4.**
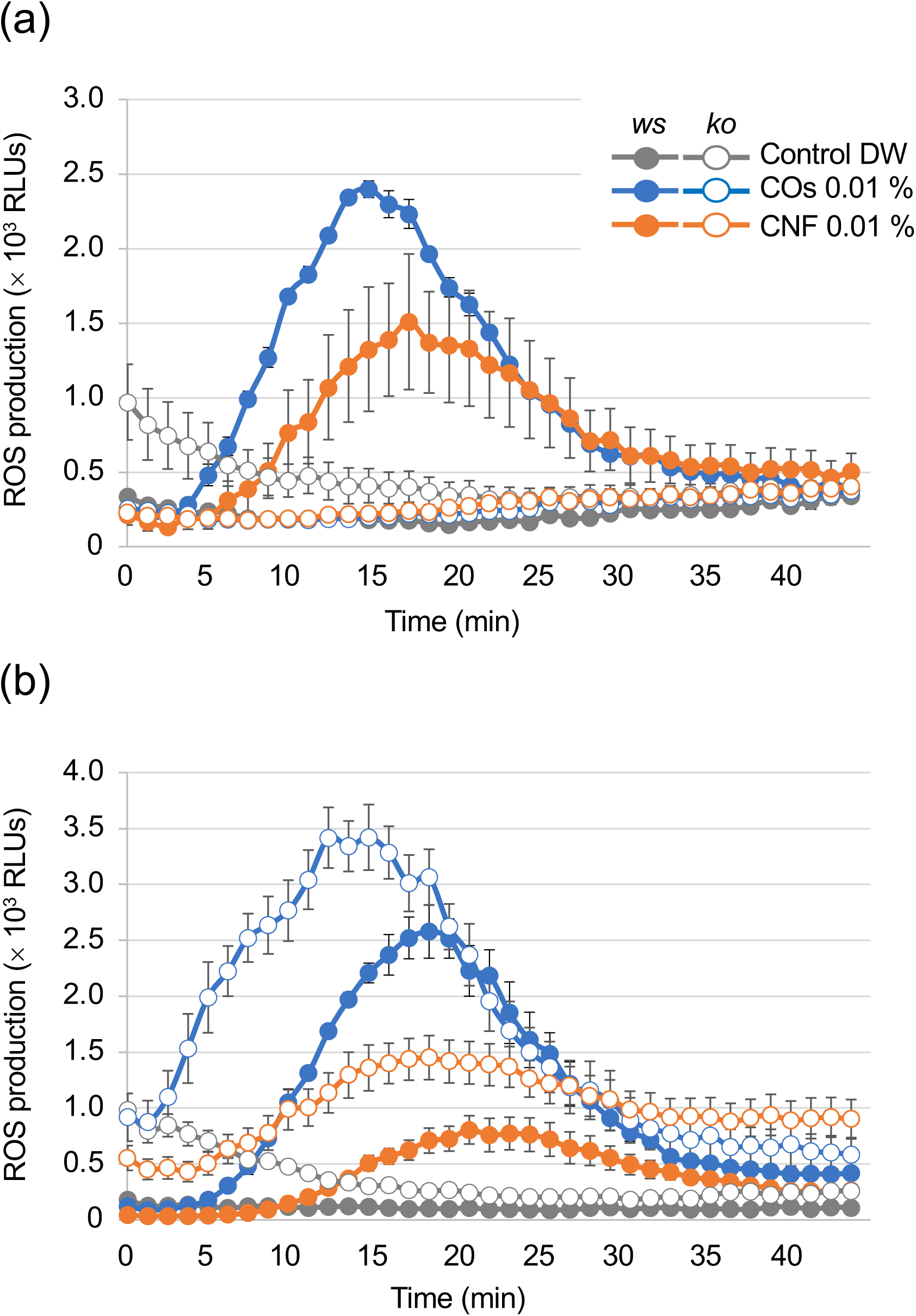
Chitins induce a local immune response in LysM-receptor mutants. (a, b) ROS production induced by chitins in rice LysM-receptor mutants. Leaf discs prepared from 3-week-old knockout (*ko*) mutants and wild-type siblings (*ws*) of *OsCERK1* (a) or *OsCEBiP* (b) grown on soil were used for ROS measurements in relative luminescence units (RLUs). Representative results from three independent experiments are shown. Error bars, SD (*n* = 3).

**Fig. 5.**
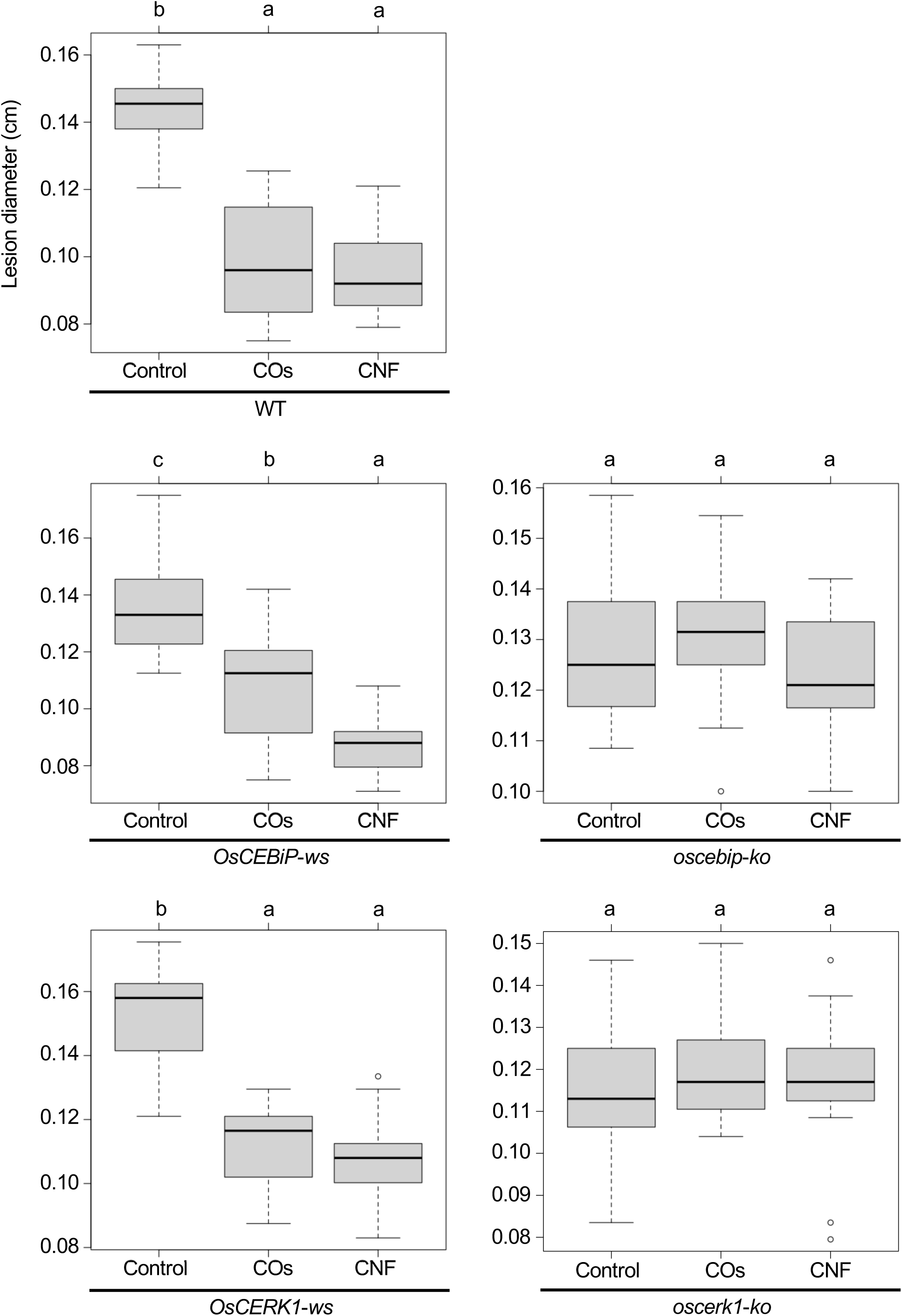
Chitins induce systemic disease resistance in LysM-receptor mutants. Lesion diameters upon *B. oryzae* inoculation in wild-type plants (WT), wild-type siblings (*ws*), and knockout (*ko*) mutants of *OsCERK1* or *OsCEBiP*, conducted as in Fig. 1. Representative results from three independent experiments are shown. Error bars, SD (*n* > 9). Different letters indicate significant differences by Tukey’s test (*p* < 0.05).

To investigate the effects of the *oscerk1* or *oscebip* mutants on chitin-induced gene expression, we performed an RNA-seq analysis on the leaves of *oscerk1* and *oscebip* mutants grown on soils mixed with chitins. Using the 297 DEGs in the wild type in response to chitin treatment as reference (Fig. **2a**), we established that the expression patterns in the *oscebip* mutant background were drastically different from those observed in the wild type and the *oscerk1* mutant (Fig. **6a**; Tables **S6**–**S9**). Next, we determined DEGs specific to the knockout mutants by comparing expression levels between control and chitin-treated seedlings and identified 1744 DEGs in *oscebip* and 1495 DEGs in *oscerk1*, of which 535 genes were common to both receptor mutants with both chitin treatments (Fig. **6b**). In fact, only 32 of the 297 chitin-induced DEGs in the wild type were differentially expressed in both knockout mutants, and 162 genes were specifically induced by chitin treatment in the wild type (Fig. **6b**). A GO enrichment analysis of these 162 genes identified the terms “mitotic cytokinesis (GO:0000281)” and “plant-type cell wall organization or biogenesis (GO:0071669)” as enriched (Fig. **6c**), which corresponded to the GO terms obtained in the DEGs downregulated by chitin supplementation (Fig. **2c**).

**Fig. 6.**
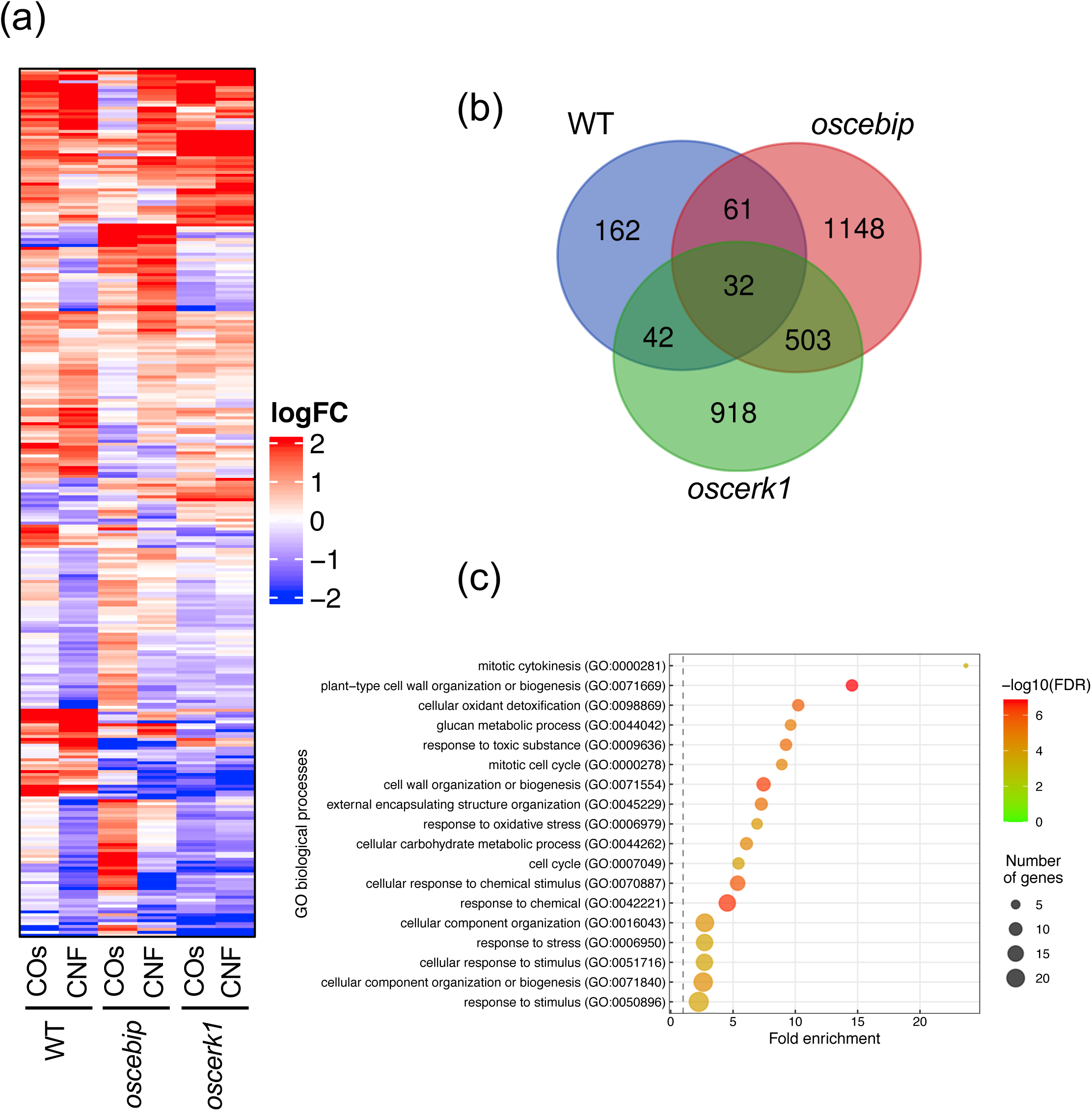
Transcriptome analysis of leaves from LysM-receptor mutants grown on chitin-supplemented soils. (a) Heatmap representation of expression levels of genes identified in the wild type (WT) as being differentially expressed in leaves upon chitin treatment and listed in Table **S2** and **S3** in the leaves of the WT and *oscebip* or *oscerk1* mutants. LogFC is shown between −2 and 2, with outside values indicated as 2 or −2. Red, upregulated; blue, downregulated. (b) Venn diagram showing the overlap between chitin-induced DEGs in the WT and genes that were DEGs in the knockout mutants (blue: in the WT, red: in *oscebip* mutants, green: in *oscerk1* mutants). (c) Results of GO enrichment analysis of the genes not differentially expressed in *oscebip* and *oscerk1* mutants defined above [162 genes; in the WT-specific group of (b)].

## Discussion

This study aimed to elucidate the molecular mechanism underlying the systemic resistance induced by chitins in rice. To this end, we used two types of chitins, COs (DP2-6) and polymeric chitin CNF, and determined their effects on systemic disease resistance, ROS production, and the transcriptome using knockout mutants of the well-characterized LysM-containing chitin receptors, OsCERK1 and OsCEBiP. Both COs and CNF induced ROS production, and we showed that OsCERK1, but not OsCEBiP, is required to elicit ROS production by chitins in rice leaves (Figs. **1a**, **4a, 4b**). Supplementation of soils with COs or CNF significantly induced systemic disease resistance against *B. oryzae* in rice leaves of wild-type plants and wild-type siblings of the knockout mutants (Figs. **1b**, **5**), while both *oscerk1* and *oscebip* mutants compromised chitin-induced systemic disease resistance (Fig. **5**). These results indicated that OsCERK1 and OsCEBiP regulate chitin-induced systemic disease resistance, although a chitin-induced local immune response in leaves likely requires another chitin-binding protein(s). When chitins are supplemented in soils, chitin perception would be expected to take place in roots and then initiate a long-distance signalling from roots to shoots to induce systemic disease resistance in leaves. Since both OsCERK1 and OsCEBiP are required for chitin-induced systemic disease resistance, these LysM receptors should function as chitin receptors in roots, but OsCEBiP is not required for ROS production in leaves (Fig. **4b**). However, the *oscebip* mutant also compromises elicitor activity in rice suspension cultured cells (Kouzai *et al*., 2014b). This discrepancy may be explained by the different materials used for analysis and ROS measurements: photosynthetic (leaves) versus non-photosynthetic (cell suspensions).

In rice leaves, COs induced ROS production more strongly than CNF (Fig. **1a**). By contrast, CNF supplementation of soils resulted in more DEGs than CO supplementation (Fig. **2**), as was previously observed with the transcriptomes of soybean (*Glycine max*) roots grown in soils mixed with COs or CNF (Kaminaka *et al*., 2020). In addition, the RNA-seq analysis of the chitin receptor mutants demonstrated that the expression patterns of DEGs in rice leaves are similar between soils supplemented with COs and CNF, with some differences as well (Fig. **6**). CNF can be degraded into oligomeric chitins by chitinase more rapidly than non-nanofibrillated chitin (Egusa *et al*., 2015). Thus, CNF treatment may be caused by these degraded forms of oligomeric chitins rather than directly by CNF. However, previous reports indicated that AtCERK1 also binds to polymeric chitin, which plays an essential role in chitin signalling (Petutschnig *et al*., 2010; Wan *et al*., 2012). Therefore, oligomeric and polymeric chitins may have specific roles in local immune responses and systemic disease resistance.

RNA-seq analysis of the leaves of rice seedlings grown on soils supplemented with chitins revealed the involvement of cell-wall biogenesis, cytokinin signalling, and regulation of phosphorylation in the systemic response induced by chitins (Fig. **2**). Cell-wall biogenesis may play a key role in chitin-induced systemic response, as this function would require the chitin receptors OsCERK1 and OsCEBiP (Fig. **6b, c**). This finding was also supported by the evidence that chitin supplementation of soils disturbs cell-wall composition, as evidenced by FT-IR spectrometry (Fig. **3a**). The leaves of chitin-treated rice seedlings showed spectra quite similar to those of the cellulose-deficient mutant *procuste 1-8* (*prc1-8*) and the pectin-deficient mutant *quasimodo 1-1* (*qua1-1*) of Arabidopsis (Mouille *et al*., 2003). We observed the same lower absorbance from 1170 to 1050 cm^−1^ in chitin-treated rice seedlings that was attributed to cellulose and xyloglucans observed in the Arabidopsis *powdery mildew resistant 5* (*pmr5*) and *pmr6* mutants, which exhibit enhanced resistance to powdery mildew (Vogel *et al*., 2004). In addition, the cellulose biosynthesis inhibitor ISX significantly induced disease resistance in leaves (Fig. **3b**). ISX targets cellulose synthase (CESA) subunits in Arabidopsis (Desprez *et al*., 2002; Scheible *et al*., 2001), which is in line with the strong reduction in the expression of genes encoding CESA or cellulose synthase-like (CSLA) among CNF-induced DEGs compared to control seedlings (Fig. **S5**). Cell-wall-derived oligosaccharides released from hemicellulose activate the immune response via OsCERK1 during infection by the fungal pathogen *Magnaporthe oryzae* in rice (Yang *et al*., 2021). Thus, damage-associated molecular pattern (DAMP)-triggered immunity caused by cell-wall-derived molecules, particularly cellulose, might be involved in chitin-induced systemic resistance in rice.

We measured a significant reduction in cytokinin levels in leaves of rice seedlings grown on CNF-supplemented soils (Table **2**). ISX treatment reduces the contents of the active cytokinin tZ as well as iP types in Arabidopsis (Gigli-Bisceglia *et al*., 2018). The expression levels of several genes encoding type-A response regulators, which regulate cytokinin signalling, were lower upon supplementation of soils with both chitins in rice leaves (Fig. **S5**). Loss of function of type-A ARR6 (Arabidopsis Response Regulator 6) induces disease resistance to *P. cucumerina* BMM and *Hyaloperonospora parasitica* Noco2 and is accompanied by an alteration in cell-wall composition (Bacete *et al*., 2020). These findings suggest that downregulation of genes involved in cytokinin signalling, which is associated with alterations of cell-wall components, participates in chitin-induced systemic disease resistance.

LysM-containing receptors perceive ligands for both immune responses and when establishing symbiosis. In rice, OsCERK1 is involved in recognizing both immune and symbiotic signals. For chitin-triggered immunity, OsCERK1 forms a receptor complex with OsCEBiP that binds to long-chain COs such as chitooctaose (CO8) (Hayafune *et al*., 2014; Shimizu *et al*., 2010). OsMYR1/OsLYK2, which directly binds to short-chain COs like chitotetraose (CO4) released by beneficial symbiont AM fungi, forms a heteromer with OsCERK1 to establish AM symbiosis (He *et al*., 2019). Disruption of OsCERK1 decreases the colonisation of AM fungi and the production of calcium spikes, whereas the *oscebip* mutant does not have any effect on symbiosis (Carotenuto *et al*., 2017; Miyata *et al*., 2014). OsMYR1 depletes OsCERK1 for OsCERK1-OsCEBiP formation and prevents immune signalling induced by CO8, while OsCEBiP inhibits OsCERK1-OsMYR1 binding in a CO8-dependent manner (Zhang *et al*., 2021). This competition between OsCERK1-OsCEBiP and OsCERK1-OsMYR1 might balance immunity and symbiosis (Zhang *et al*., 2021). Since the systemic induction of disease resistance by chitins appears similar to what takes place during ISR caused by beneficial fungi, chitin-induced systemic disease resistance may employ the recognition mechanism for chitins involved in local immune response via OsCERK1-OsCEBiP but not the OsCERK1-OsMYR1 receptor complex participating in AM symbiosis. In addition, the expression of DEGs upregulated by chitins in leaves displayed a strong positive correlation with the genes induced by BTH, an SA analogue that induces defence priming (Shimono *et al*., 2007) (Table **1**). Our previous study reported that CNF supplementation of soils induces defence priming in cabbage and strawberry (Parada *et al*., 2018). Taken together with the evidence that fungal ISR caused by *L. bicolor* in Arabidopsis occurs via AtCERK1 (Vishwanathan *et al*., 2020), chitin-induced systemic disease resistance may mimic ISR induced by plant growth–promoting fungi.

In summary, chitins supplemented into soils systemically induce disease resistance against the fungal pathogen *B. oryzae* via recognition of chitins by the LysM receptors OsCERK1 and OsCEBiP in rice. Cell-wall biogenesis and cytokinin signalling are perturbed as a systemic response in leaves, and defence priming-related genes and phosphorylation-related genes are upregulated. These effects, together with another unknown function, eventually induce disease resistance (Fig. **7**). This study uncovers the molecular basis underlying chitin-induced systemic disease resistance. These findings may also contribute to elucidating the molecular basis of ISR, which is not well understood, and provides support for the application of chitins as a promising material in agriculture to confer disease resistance. However, it remains unknown how plants systemically induce disease resistance in response to chitins. In addition, both oligomeric and polymeric chitins caused similar effects on chitin-induced systemic disease resistance but differently affected local immune responses and global gene expression in leaves. Thus, it will be essential to expand our knowledge regarding chitin-induced disease resistance in rice, for example, by identifying the molecules involved in long-distance signalling and those derived from cell walls and by confirming the direct perception of CNF by LysM receptors.

**Fig. 7.**
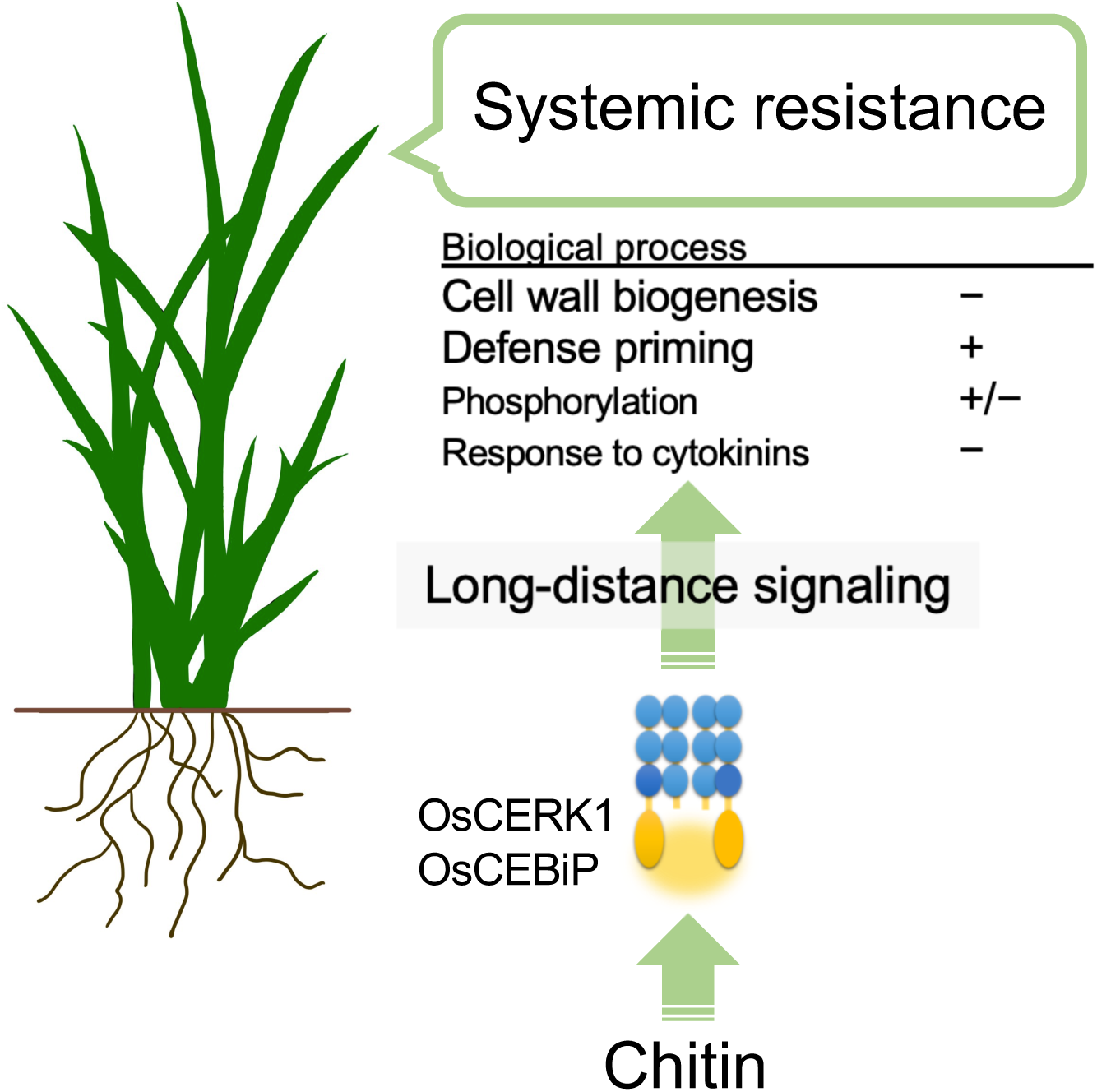
Hypothetical model of chitin-induced systemic disease resistance in rice. Chitins supplemented in the soil induce disease resistance in leaves against the fungal pathogen *B. oryzae*. Chitins are first recognized by the LysM receptors OsCERK1 and OsCEBiP in roots. Then, long-distance signalling initiated in the roots perturbs cell-wall biogenesis and cytokinin signalling and upregulates defence priming-related genes and phosphorylation-related genes in leaves, inducing disease resistance.

## Acknowledgements

We would like to thank Dr. Makoto Ueno (Shimane University) and Dr. Atsushi Ishihara (Tottori University) for providing the *B. oryzae* strain. We also thank Ms. Mei Yokomizo for technical assistance. This work was supported by JSPS KAKENHI Grant-in-Aid for Scientific Research (B) (Grant no. 19KT0010) and Takahashi Industrial and Economic Research Foundation.

## Author Contributions

YN, SI, AM, and HK conceived and designed the experiments; TM, KH, KN, SM, ME, YK, MS, and AM performed the experiments; TM, KH, and AM analysed the sequencing data; TM, ME, YN, SI, AM and HK wrote the manuscript. All authors approved the final manuscript.

## Data availability

The raw read data for RNA-seq were deposited in the DNA Data Bank of Japan under the accession number DRA012267.

## Supporting information

**Fig. S1.** Effects of chitin supplementation of the soil on the growth of rice seedlings.

**Fig. S2.** Venn diagram showing the overlap between the number of DEGs in rice leaves treated with COs or CNF.

**Fig. S3.** BTH-induced genes are also upregulated in response to chitin treatments.

**Fig. S4.** Results of RNA-seq analysis of chitin-treated rice roots.

**Fig. S5.** Expression levels of cell-wall- and cytokinin-related genes.

**Table S1.** Summary of RNA-seq analysis.

**Table S2.** DEGs in wild-type rice leaves induced by supplementation of soils with COs.

**Table S3.** DEGs in wild-type rice leaves induced by supplementation of soils with CNF.

**Table S4.** DEGs in wild-type rice roots induced by supplementation of soils with COs.

**Table S5.** DEGs in wild-type rice roots induced by supplementation of soils with CNF.

**Table S6.** DEGs in leaves of the *oscebip* knockout mutant induced by supplementation of soils with COs.

**Table S7.** DEGs in leaves of the *oscebip* knockout mutant induced by supplementation of soils with CNF.

**Table S8.** DEGs in leaves of the *oscerk1* knockout mutant induced by supplementation of soils with COs.

**Table S9.** DEGs in leaves of the *oscerk1* knockout mutant induced by supplementation of soils with CNF.

**Table S10.** DEGs in leaves of the wild type induced by supplementation of soils with chitin that are not included in the DEGs induced by chitin treatment in the *oscebip* or *oscerk1* mutants.

