## Supplemental Figure 1-5 for "Chitin-induced systemic disease resistance in rice requires both OsCERK1 and OsCEBiP and is mediated via perturbation of cell-wall biogenesis in leaves"

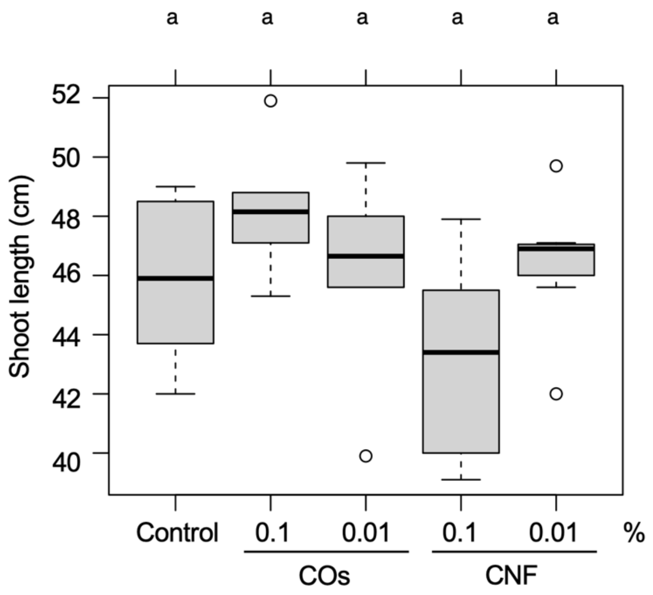


**Fig. S1. Effects of chitin supplementation of the soil on the growth of rice seedlings.** The seedlings were grown on soils and treated with distilled water (Control), chitin oligomers (COs), or chitin nanofiber (CNF) at indicated final concentration (w/v). Error bars, SD ($n$ = 6). Differences in each sample were not significant, according to Tukey's test ($p < 0.05$).

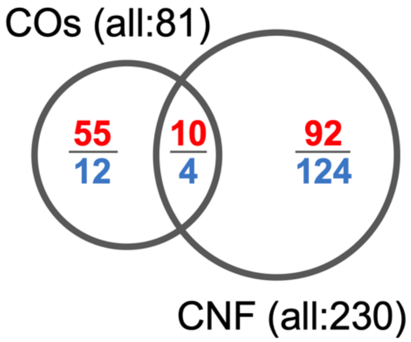


**Fig. S2. Venn diagram showing the overlap between the number of DEGs in rice leaves treated with COs or CNF.** The number indicates upregulated (Red) or downregulated (Blue) genes.

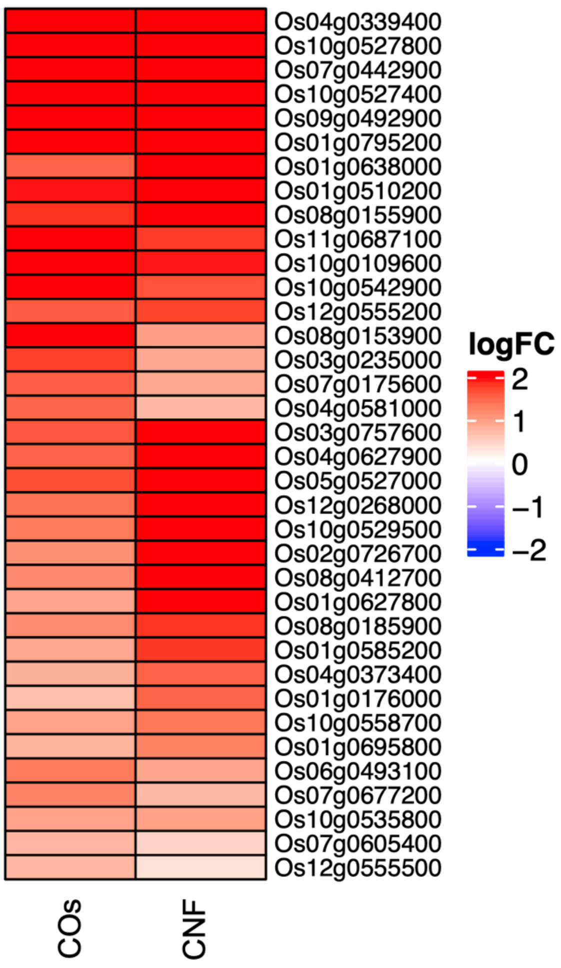


**Fig. S3. BTH-induced genes are also upregulated in response to chitin treatments.** Expression levels of genes upregulated by chitin treatments correlated with BTH-induced genes listed in Table 1 are visualized by heatmaps. LogFC is shown between –2 and 2, with outside values indicated as 2 or –2. Red, upregulated; blue, downregulated.

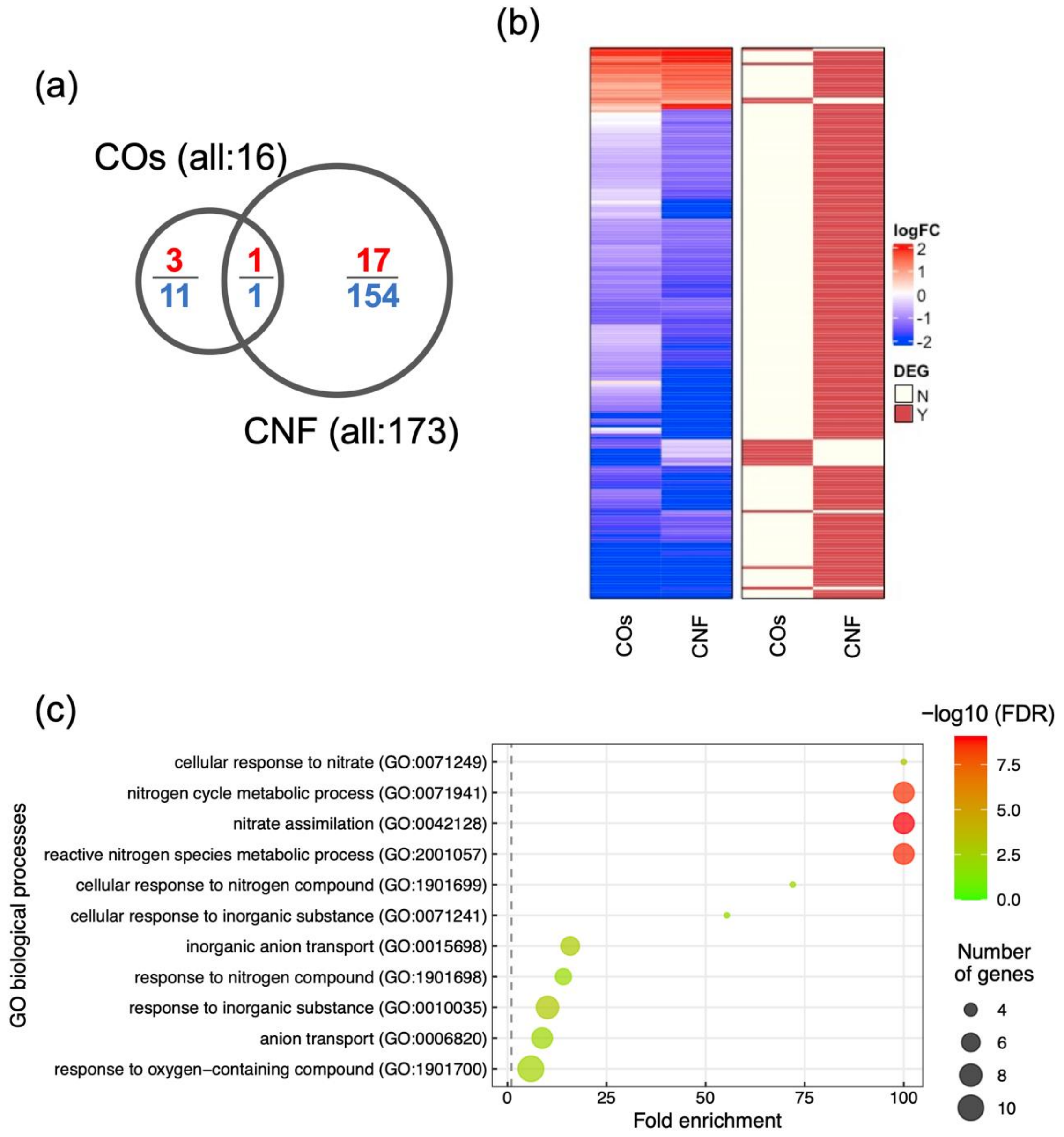


**Fig. S4. Results of RNA-seq analysis of chitin-treated rice roots.** (a) Venn diagram for the number and grouping of differentially expressed genes (DEGs). The number indicates upregulated (Red) or downregulated (Blue) genes. (b) Expression levels of each gene are visualized by heatmaps (left). LogFC is shown between −2 and 2, with outside values indicated as 2 or −2. Red, upregulated genes; blue, downregulated genes. DEGs in each treatment are indicated on the right. Red, DEG; ivory, not differentially expressed. (c) Results of Gene Ontology (GO) enrichment analysis summarized as plot data.

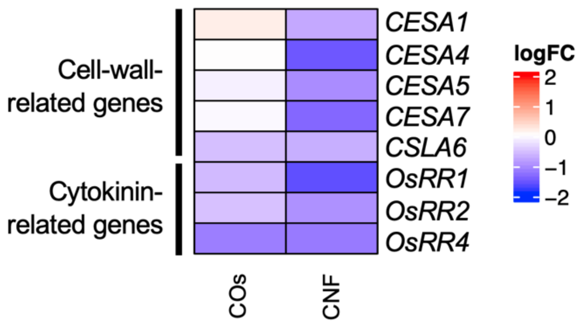


**Fig. S5. Expression levels of cell-wall- and cytokinin-related genes.** Expression levels of *CESA* (*cellulose synthase* ), *CELA* (*cellulose synthase-like A* ), and *RR* (*response regulator* ) genes in leaves of rice plants grown on soils mixed with 0.01% (w/v) COs and CNF. Red, upregulated; blue, downregulated.
